# Global population structure of a unicellular marine predator

**DOI:** 10.1101/2024.11.27.625609

**Authors:** Francisco Latorre, Olivier Jaillon, Michael E. Sieracki, Corinne Cruaud, Ramon Massana, Ramiro Logares

## Abstract

Unicellular predators, especially marine heterotrophic flagellates (HFs), are fundamental to marine food webs, facilitating the flow of nutrients from bacteria and the smallest primary producers to upper trophic levels. Comprehending their distributions and diversity is essential for understanding the functioning of marine ecosystems. The uncultured MArine STramenopiles clade 4 (MAST-4) constitutes a crucial HF member of the ocean microbiome with a predatory (bacterivorous) lifestyle. Temperature gradients have influenced the evolution and biogeography of MAST-4 species. Yet, similarly to most uncultured microbes, there is limited information on the population diversity and structure within MAST-4 species. This information is key for better comprehending the ecological roles and adaptations of different MAST-4 species to specific ocean niches. Here, we investigate the population diversity and structure on the global scale of the MAST-4 species A, B, C, and E by combining metagenomics and single-cell genomics data from the *Tara Oceans* expedition. We found substantial population divergence in the abundant MAST-4A and C species, while lower levels of divergence were observed in species B and E. Temperature and salinity were the main drivers structuring the populations within the four MAST-4 species. The analysis of positively selected gene clusters within each MAST-4 species revealed genomic areas likely the basis of population adaptation to different niches. Overall, our results contribute to a better comprehension of the population diversity and structure of four species belonging to an important group of unicellular predators, offering insights into their ecological roles and adaptations in the global ocean. The results also contribute to understanding microbial populations, a dimension of diversity that remains poorly known for most wild species and is pivotal in global change.

## INTRODUCTION

Heterotrophic unicellular eukaryotes populate all aquatic ecosystems, including sediments and other organisms, from the surface to the dark ocean^1,2^. In the ocean, heterotrophic flagellates (HF) represent around 20% of the total eukaryotic cells in the sunlight zone^3^. HFs are crucial in marine food webs by channeling nutrients and energy from bacteria and the smallest primary producers to upper trophic levels. These organisms are active grazers, important agents in regulating prokaryotic and small-eukaryotic abundances in the plankton^4,5^. Studies during the last three decades have shown that HFs comprise evolutionary divergent organisms affiliating with all major eukaryotic supergroups^6–8^. Yet, HFs are often analyzed as a single functional group, overlooking the organismal and evolutionary differences of the species within this key assemblage.

Marine plankton, including HFs, can travel thousands of kilometers while being carried by currents. During their drift, HF cells from the same species may encounter diverse habitats, from polar to tropical waters and sunlit surfaces to the deep, dark ocean, each presenting unique ecological conditions and challenges. Such environmental heterogeneity can exert disruptive selection and may lead to local adaptation in different populations^9^. The genomic variation between intraspecific individuals may evidence such adaptations. These may be represented by accessory genes, duplicated genes, Single Nucleotide Variants (SNVs), Multiple-Nucleotide Variants (MNVs), Insertions, Deletions, or structural variants^10,11^.

Population-level studies have assessed the adaptation and other evolutionary processes in macrofauna^12–14^ and a number of microorganisms, primarily prokaryotes^15–17^ and marine phytoplankton^18^. Nevertheless, little is known about the population diversity and structure of most wild microbes, particularly HFs^11^. Addressing this knowledge gap is essential for advancing our understanding of the ocean microbiome and its intricate connection to ecosystem function. Today, metaomic techniques, such as metagenomics, can help us to investigate this uncharted dimension of diversity^11^. Given the large population sizes and fast reproduction rates of HFs, their populations are expected to harbor substantial genetic diversity. This is because larger populations can sustain more alleles, reducing the effects of genetic drift. At the same time, fast reproduction rates increase the opportunities for mutation and recombination, further contributing to genetic variability^11^. This diversity is likely organized into locally, regionally, or temporally adapted populations, reflecting environmental heterogeneity and biotic interactions.

Among HFs, Marine Stramenopiles (MASTs), with cell sizes ranging between 2-5 µm, are the most frequent in the surface ocean^19–22^. Most are uncultured^22–24^ and can account for up to 50% of the microeukaryote abundance in metabarcoding analyses in specific locations^25^. So far, 18 MAST groups have been recognized, branching from different basal areas of the stramenopile phylogeny^26^. MASTs can inhabit oceanic waters from the surface (MAST-1, -3, -4, and -7)^19,23,26,27^ to deeper layers (MAST-23)^26^, from tropical (MAST-1 and -4)^27,28^ to polar areas (MAST-2)^25,29^, and also populate freshwater environments (MAST-2 and -12)^26,30^. MASTs are essential in food webs as active grazers on bacteria and tiny algae^24,28,31,32^. Some have been described in symbiotic-like forms with unicellular algae, either diatoms or cyanobacteria (MAST-3)^33^, suggesting the possibility for mixotrophy in this group.

Among the MASTs, the MAST-4 clade shows a relatively low species diversity, worldwide distribution, and high relative abundance in surface marine waters compared to other HFs (∼9% of all HFs)^22^. As a result, MAST-4 is becoming a model organism for studying HFs’ ecology in the ocean. The biogeography of MAST-4 lineages has been partially elucidated, and species distributions appear primarily driven by abiotic and biotic environmental heterogeneity^34^. This suggests that their diversification in the surface ocean has been promoted by adaptation^27^. In a previous study^27^, we found that MAST-4B and C co-occur in tropical waters, having a limited overlap with the distribution of MAST-4A, which predominantly inhabits subtropical waters. Temperature and, to a lesser extent, salinity were identified as the main drivers shaping the biogeography of MAST-4 species. Genomic analyses further suggested that prey selection and competition for prey contributed to the diversification of MAST-4. Therefore, biotic interactions seem to have also played a significant role in shaping the biogeography of MAST-4 species.

While recent data on the biogeography of MAST-4 species have provided promising insights, a substantial knowledge gap remains regarding distinct populations and their distribution patterns across the global ocean. This was primarily due to challenges obtaining genomic data from these uncultured HFs. Today, single-cell genomics (SCG) combined with metagenomics enables the study of uncultured marine protists’ populations through "metagenome-based population genomics." Specifically, SCGs has demonstrated its capability to yield partial and sometimes nearly complete genomes of uncultured protists, as evidenced by its application to MAST-4 species^24,27,35–37^. Then, the population-level variation over spatiotemporal scales can be inferred using metagenomic data^11^. This is achieved by mapping short metagenomic reads from environmental DNA against protistan genomes and identifying Single-Nucleotide Variants (SNVs), small insertions or deletions (indels), and population-specific gene losses or insertions.

The metagenome-based population genomics approach has been used to investigate aquatic prokaryotes’ population differentiation, diversity, and adaptation. In contrast, few studies have used this approach to investigate protist populations^11^. In one study, Leconte and colleagues examined the picophytoplankton *Bathycoccus* using a reference genome isolated from the Western Mediterranean Sea^17^. They assessed broad patterns of population-level variation using metagenomes from surface waters and deep chlorophyll maximum layers collected during the *Tara Oceans* campaign, focusing on the 0.8–5 μm size fraction. Of the 162 original metagenomes from the *Tara Oceans* dataset, only 27 from various geographic locations and ocean basins had sufficient reference genome coverage for downstream analyses. When the 27 metagenomes were compared based on the SNVs identified in the reference genome, a distinct separation emerged between those from the Arctic and temperate regions. Additionally, Arctic populations were separated from Austral ones, and a positive correlation was observed between population differentiation and temperature differences^17^, suggesting that temperature plays a significant role in shaping the population genomic variation of *Bathycoccus*. Furthermore, 2,742 SNVs and 13 single amino acid variants (SAAVs) were identified, differentiating temperate from cold-water *Bathycoccus* populations. Comparisons of protein variants from mesophilic and psychrophilic populations provided insights into structural changes that may drive adaptation to different thermal niches by impacting the protein’s functional and physical properties^17^.

In another study, Da Silva and colleagues explored the genomic differentiation among three species of pico-phytoplankton in the Mediterranean Sea: *Bathycoccus prasinos*, *Pelagomonas calceolata*, and *Phaeocystis cordata*^38^. Metagenomic reads from *Tara Oceans* stations in the Mediterranean Sea were mapped to reference genomes for *B. prasinos* and to transcriptomes for *P. calceolata* and *P. cordata* obtained from the Mediterranean Sea or other regions. *B. prasinos* exhibited higher population differentiation in the Mediterranean Sea than *P. calceolata* and *P. cordata*. Moreover, environmental selection appeared to influence the population diversity of *B. prasinos*, whereas *P. cordata* populations seemed to be shaped primarily by geographic distance^38^. Thus, populations of different protist species within the same functional group and with similar morphologies can exhibit substantial variation in their levels of differentiation. Distinct processes, such as natural selection or dispersal may drive this variation. Although still in its early stages, metagenome-based population genomics offers significant potential to reveal uncultured protists’ population characteristics and structure^11^.

Here, we investigate the global population genomics of four MAST-4 species (A, B, C, and E) using genomes obtained via single-cell genomics^27^ and surface ocean metagenomes from the *Tara Oceans* expedition. We aimed to determine the amount of genetic diversity within the analyzed MAST-4 species in the global surface ocean. We also analyze the presence of population structure and identify the possible environmental or geographic drivers. Furthermore, we search for genes or genomic regions in different populations that may be the basis of their adaptation to diverse niches. The results expand our understanding of the population diversity, structure, and adaptive diversification of a critical marine unicellular predator, being relevant for comprehending the role of HFs in the functioning of the ocean microbiome.

## METHODS

### Genome reconstruction using Single Amplified Genomes

Genomes from MAST-4 species A/B/C/E were reconstructed in a previous work^27^ after co-assembling multiple Single Amplified Genomes (SAGs) from plankton samples collected during the circumglobal *Tara Oceans* expedition^21^. A total of 69 SAGs (over 424.1 Gb of sequencing data) were selected and processed to generate a single co-assembled genome for each species. For each co-assembly, estimations of genome recovery were calculated with BUSCO v3 (Benchmarking Universal Single-Copy Orthologs)^39^ using the eukaryote_odb9 dataset. The identified orthologs were then used as a training set for AUGUSTUS 3.2.3^40^ to predict genes. Functional annotation was performed using different databases [KEGG (Release 2015-10-12)^41,42^, eggNOG v4.5^43^, and CAZy database from dbCAN v6^44,45^] with HMMER 3.1b2^46^ and BLAST 2.2.28+^47^. More details on the assembly and cleaning processes of the MAST-4 genomes are described in Latorre *et al.*^27^

### The abundance of MAST-4 genomes in the ocean

We determined the abundance of the four studied MAST-4 genomes in the global ocean. To achieve that, we mapped 111 metagenomes from *Tara Oceans* (82 surface ocean stations encompassing the 0.8 – 5 μm size fraction) against the whole genome of each MAST-4 species (**Supplementary dataset S1 and dataset S2**). BAM files for each station and genome were generated with BWA 0.7.17-r1188^48^, and only reads with identity > 95% and an alignment coverage > 80% were kept. Abundance values estimated as RPKM (reads mapped per kilobase of the genome, per million mapped read) were calculated from the BAM files for each station using CoverM v0.7.0^49^ under default parameters. Genomic Horizontal Coverage, defined as the percentage of the genome covered by at least one filtered read, was calculated using the number of covered genome bases provided by CoverM. We only considered samples (metagenomes) covering at least 25% of the genome for in-depth analyses.

### Genetic diversity and divergence of MAST-4 in the open ocean

To assess the genetic diversity and divergence within MAST-4 species in the global ocean, we analyzed Single Nucleotide Variants (SNVs) and small insertions or deletions (indels) across different ocean regions. For each MAST-4 genome, 82 BAM files (one per station) were merged into a single BAM file using the *Samtools 1.8 merge* function. Merged BAM files were input to *Freebayes* v1.3.1^50^ to perform variant calling, with a ploidy set to 1 (-p 1) and the minimum number of observations to support an alternate allele set to 4 (-C 4). The resulting variant call files (VCF) were used as input to a) *SnpEff 5.0e* (build 2021-03-09)^51^ with default parameters to annotate and predict the effects of genetic variants on MAST-4 genomes, genes, and proteins and b) *POGENOM v.0.8.3*^52^, with minimum coverage for a locus set to 10 (--min_count 10) and minimum number of stations that a locus needs to be present to 4 (--min_found 4). This analysis facilitated the computation of nucleotide diversity (π), defined as the average number of nucleotide differences per site between any two sequence reads chosen randomly from the sample population, and the Fixation index (FST), which measures the proportion of genetic diversity due to allele frequency differences among all the possible pairwise population comparisons between stations.

Following Hartl and Clark^53^, we established four groups of genetic differentiation based on FST pairwise values: FST < 0.05, little genetic divergence; 0.05 < FST < 0.15, moderate; 0.15 < FST < 0.25, high; FST > 0.25, very high. The amount of genetic differentiation (FST values) explained by selected environmental variables (temperature and salinity) was analyzed with PERMANOVA (*adonis* function in the *vegan* R-package) using environmental variables with Z-score normalization. Station 11 was removed from these analyses due to missing data.

### Calculation of dN/dS ratios

Potential adaptive evolution in MAST-4 coding sequences was analyzed using the ratio of non-synonymous vs. synonymous substitutions (dN/dS). Overall, it is considered that a dN/dS > 1 implies positive or diversifying selection, dN/dS < 1 purifying or negative selection, and dN/dS = 1 neutral evolution^54^. Here, we calculated the dN/dS ratios per gene and station following Nei and Gojorobi^55^ and Morelli et al., 2013^56^ using a *Python script* developed *ad hoc* (available at *https://github.com/franlat/mast4_dnds.git*). In this approach, we compensated for the biases introduced by sequencing data coverage differences by normalizing the synonymous and non-synonymous counts using codon coverage.

## RESULTS

### Variant detection and annotation in MAST-4

For each MAST-4 species, genomic variants were classified into Single-Nucleotide Variants (SNVs), Multiple-Nucleotide Variants (MNVs), insertions and deletions (INDELs), and all the possible combinations (MIXED). A total of 864,009/131,091/668,613/137,357 genomic variants (18.2/4.5/14.0/4.5 variants per kb of genome) were predicted for MAST-4A/B/C/E, respectively. On average, 87% of the variants in each MAST-4 were SNVs, while the other 13% was distributed among MNVs, INDELs, and MIXED. Considering that one variant can have more than one effect on different genes (*e.g.,* an SNV in the downstream area of gene A can also be part of the upstream area of gene B), a total of 2,644,001/437,930/2,196,738/446,874 effects (3.06/3.34/3.29/3.25 effects per variant) were annotated for MAST-4A/B/C/E. On average, 78.4% were located in non-transcribed areas of the genome, 20.4% in coding regions, and 1.2% in non-coding regions (**Table S1**). Variants in the coding areas were classified into missense or silent variants, depending on whether they change the resulting amino acid sequence or not, and nonsense variants, if they truncate the resulting protein by introducing a stop codon. MAST-4A and E displayed an average of ∼ 45% missense and ∼ 55% silent variants, while MAST-4B and C showed an average of ∼ 32% and ∼ 68% respectively. Less than 1% of the variants’ effects were assigned as nonsense.

The effects of the variants were assigned to impact categories: HIGH, when the variant is assumed to have a disruptive impact on the protein (truncation or loss of function); MODERATE, when a variant can potentially change the protein effectiveness; LOW, when the variant is unlikely to shift protein functionality; and MODIFIER, for non-coding variants or variants for which it is difficult to predict impact. The impact of variants on MAST-4A, B, and C were, on average, proportionally similar for each category: 0% - HIGH, 13.4% - MODERATE, 8.6% - LOW, and 78.0% - MODIFIER. In turn, MAST-4E had proportionally more effects identified as MODIFIER and less as LOW in comparison to the other MAST-4 species (0.1% - HIGH, 8.6% - MODERATE, 7.2% - LOW, and 84.1% - MODIFIER) (**Table S1**).

### Genetic divergence of MAST-4 populations

For each MAST-4 genome, we calculated the fixation index (FST) using the allele frequency differences between pairs of *Tara* Ocean stations (samples). The FST measures the genomic difference between the populations in two samples, with values ranging from zero to one. A zero value means no genetic difference between populations in both samples (i.e., the same population is present in the samples), while higher values indicate increasing population differentiation^57^. We delineated four groups of genomic differentiation based on pairwise FST values, following Hartl and Clark^53^: FST < 0.05 - Little genetic divergence, 0.05 < FST < 0.15 – Moderate, 0.15 < FST < 0.25 – High, FST > 0.25 - Very high. We only considered FST values from those stations (82 total) featuring metagenomes mapped to at least 25% of a given MAST-4 genome (horizontal coverage). Thus, 50 stations were analyzed for MAST-4A, 11 for MAST-4B, 40 for MAST-4C, and 16 for MAST-4E. Only MAST-4 A and C showed FST values above 0.25, with a maximum FST value of 0.56 and 0.54, respectively (**Figure 1**). This points to very high genetic divergence between specific MAST-4A and C populations. In contrast, MAST-4B and E always displayed FST values under 0.25, with a maximum FST value of 0.14 and 0.21, respectively. All four MAST-4 species exhibited FST distance peaks at the 0.05 – 0.15 range (moderate genetic divergence) (**Figure 1**). These analyses demonstrate a substantial population divergence within the most abundant MAST-4 species in the global ocean, A and C.

**Figure 1.**
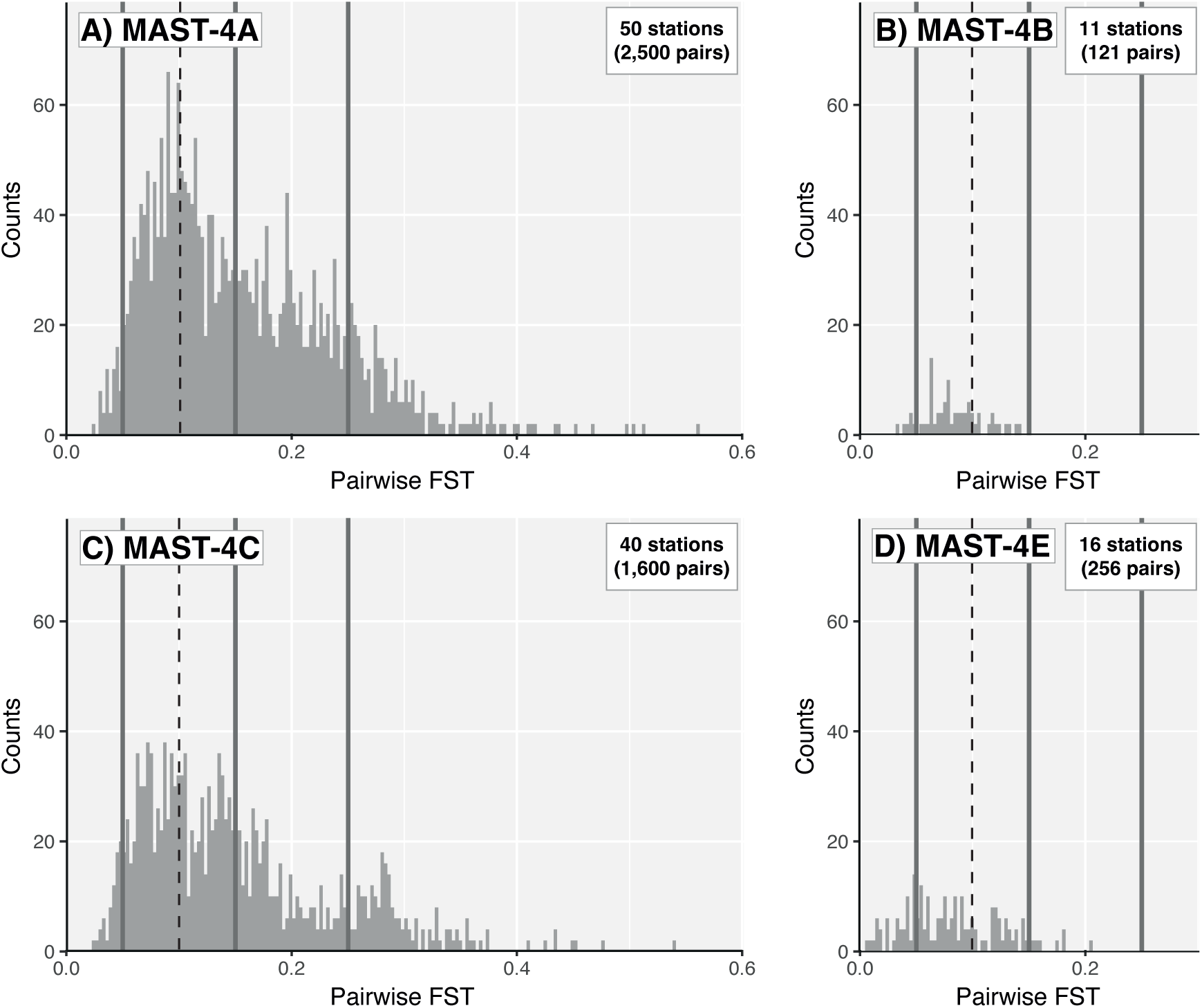
Distribution of Fixation Index (FST) values for the MAST-4 species A, B, C & E considering samples from the global ocean. Histograms show all FST values among *Tara Oceans* stations (featuring horizontal coverage ≥ 25%) for A) MAST-4A, B) MAST-4B, C) MAST-4C, and D) MAST-4E. The four MAST-4 species had FST distance peaks at the 0.05 – 0.15 range (dashed vertical line at 0.10). The solid vertical lines indicate the used differentiation categories: FST < 0.05 - Little genetic divergence, < FST < 0.15 – Moderate, 0.15 < FST < 0.25 – High, FST > 0.25 - Very high.

### Environmental heterogeneity, genetic divergence, and diversity

Temperature and salinity were the most relevant in explaining the population-level differentiation within MAST-4 A, B, C & E (*i.e.*, FST values among *Tara Oceans* stations). Temperature emerged as the predominant factor potentially influencing population differentiation in MAST-4B, C, and E, accounting for 37%, 20%, and 60% of their respective FST variance in the global ocean (PERMANOVA, p-value < 0.001). In turn, salinity stood out for species A as the primary driver, explaining 30% of the population-level differentiation, while temperature contributed to 13% (p-value < 0.001). Yet, when removing the Mediterranean stations from the PERMANOVA analysis, salinity’s importance was reduced to 13%, and temperature increased to 20%. This is coherent, as the salinity of the Mediterranean Sea stations (mean = 38.0, SD = 0.6) was higher than that of samples from other basins (mean = 35.4, SD = 1.2; Supplementary Dataset S2).

In surface waters of the open ocean, MAST-4 A and C exhibited distinct clusters, indicative of different populations, when stations were aggregated according to FST values. These clusters became apparent upon applying a 0.15 threshold for average FST distances in the UPGMA dendrograms (**Figure 2**). MAST-4A displayed a total of four clusters or potential populations. In subtropical waters, it featured one cluster in the Mediterranean Sea, designated as A1 (10 stations), and a second cluster covering other subtropical locations, labeled A2 (32 stations) (**Figure 2A**). A third potential population, A3, was identified at station 65 near the South African coast. The last cluster, A4 (7 stations), was found in tropical stations, where MAST-4A exhibited low abundance (**Figure 2A**). MAST-4C also displayed four clusters: one in the Mediterranean Sea (4 stations), denoted as C1; two in tropical waters, where MAST-4C is most prevalent, including C3 in the Arabian Sea (Indian Ocean; 5 stations); and C4 in the remaining tropical stations (30 stations) (**Figure 2C**). Additionally, MAST-4C featured one potential population, C2, exclusively found at station 65 along the South African coast.

**Figure 2.**
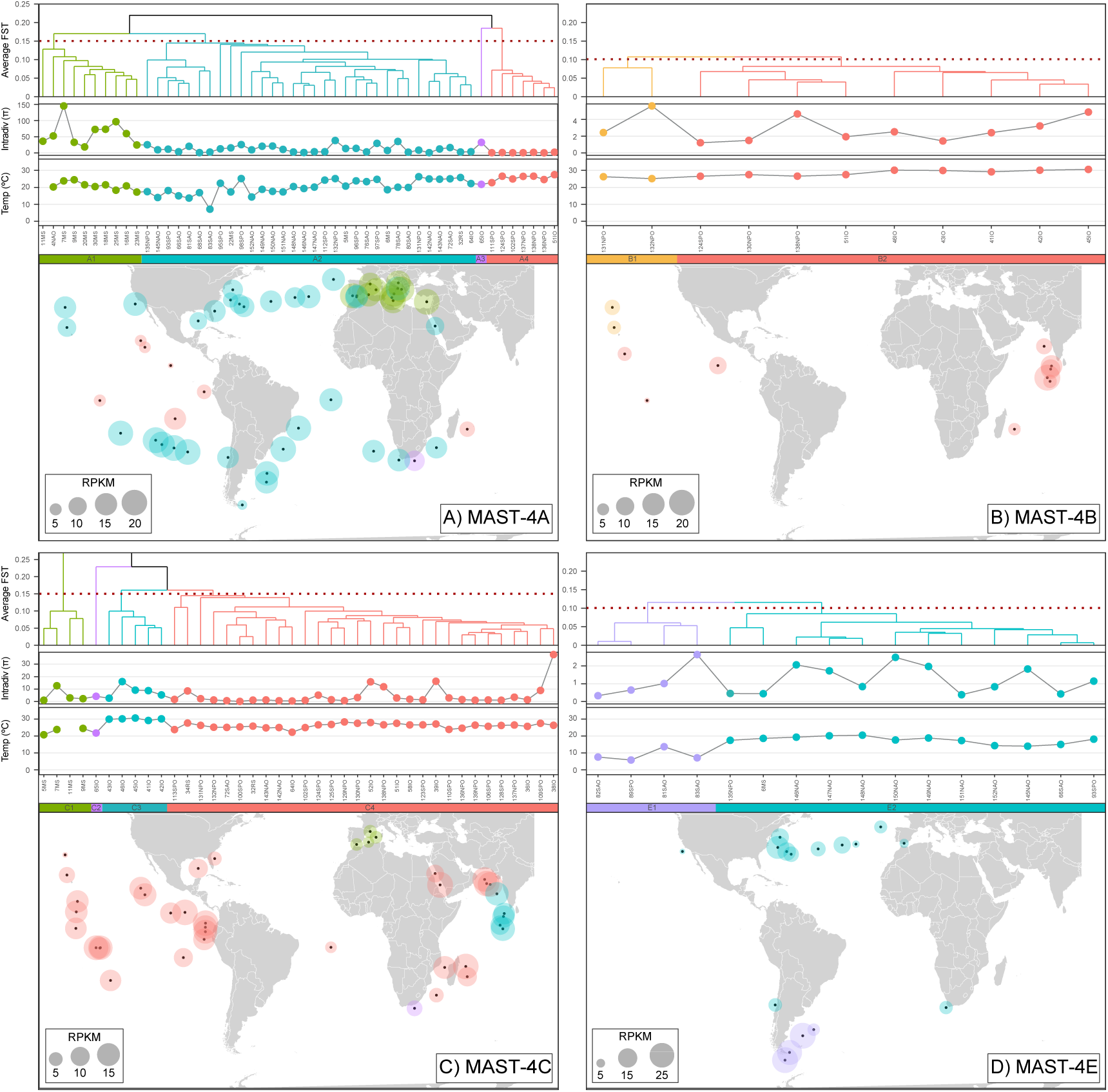
Potential populations of MAST-4A/B/C/E. Clustering of *Tara Oceans* stations (metagenomes) based on FST values for A) MAST-4A, B) MAST-4B, C) MAST-4C, and D) MAST-4E. For each species, a UPGMA dendrogram based on FST values is shown for metagenomes (stations) that covered at least 25% of each reference genome, along with the corresponding surface water temperature (important environmental driver) and the nucleotide diversity (π). The colors in the dendrograms, temperature sub-panels, nucleotide diversity sub-panels, bubbles, and those in the horizontal bars in panels A and C indicate clusters (putative populations) delineated using an FST = 0.15 threshold. Colors in panels B and E indicate clusters (putative populations or subpopulations) based on an FST = 0.10 threshold. Each population or subpopulation is identified with a letter and number in the colored horizontal bars. Bubble size represents normalized species abundances (RPKM; Reads per kilobase per million mapped reads) in a given station. Station name tags include the *Tara Oceans* station number and an acronym of the ocean region to which they belong (MS – Mediterranean Sea; RS – Red Sea; IO – Indian Ocean; SAO – South Atlantic Ocean; SO – Southern Ocean; SPO – South Pacific Ocean; NPO – North Pacific Ocean; NAO – North Atlantic Ocean).

In contrast to MAST-4A and C, MAST-4B and E each exhibited a single potential population across the surface global ocean when considering an FST threshold of 0.15. MAST-4B was more prevalent in tropical waters (**Figure 2B**), and MAST-4E was more abundant in subtropical and sub-polar waters, typically near the coast, except for specific locations in the North Atlantic Ocean (**Figure 2D**). When we lowered the FST threshold to 0.10, two potential populations or subpopulations emerged for MAST-4B and E. The putative population B1 was present in the North Pacific Ocean (2 stations), and B2 was present in the North and South Pacific Oceans, as well as in the Indian Ocean (9 stations) (**Figure 2B**). For MAST-4E, population E1 encompassed stations in the South Atlantic and Pacific Ocean (4 stations), while E2 included subtropical locations in the Mediterranean Sea, Atlantic, and Pacific Oceans (12 stations). Detailed information for each MAST-4 population, including average temperature and salinity, is shown in **Table S2**.

The analyses of nucleotide diversity did not display clear trends for the stations that were investigated (**Figure 2**), except for MAST-4A’s Mediterranean population A1, which showed a higher nucleotide diversity compared to the rest of the populations (p-value < 0.01; Wilcoxon test) (**Figure 2A**). Although less evident, MAST-4C’s C3 population in the Indian Ocean also exhibited a slightly higher nucleotide diversity on average than the rest (p-value = 0.027; Wilcoxon test). These findings suggest a higher level of variation at the nucleotide level in the individuals within these populations.

### Detecting population adaptation

When evaluating the population-level coding-gene variation across stations, the ratio between non-synonymous and synonymous mutations in coding regions (dN/dS) can provide insights into the action of natural selection. Specifically, dN/dS < 1 points to purifying or negative selection, dN/dS = 1 neutral evolution, and dN/dS > 1 positive or diversifying selection. Here, we focused on those genes featuring an average dN/dS > 1 as they could represent the basis of population differential adaptation. Genes with average dN/dS > 1 are expected in populations different from the reference MAST-4 genome. The MAST-4A populations A1, A2, A3, and A4 (**Figure 2A**), delineated with an FST=0.15 threshold, displayed mean positive selection (dN/dS > 1) in 581, 189, 679, and 102 genes, respectively. Of these, 388 (A1), 36 (A2), 504 (A3), and 52 (A4) genes were positively selected exclusively in these populations (*i.e.*, average dN/dS > 1 within a specific population only). Similarly, the MAST-4C populations C1, C2, C3, and C4 (**Figure 2C**) showed positive selection in 93, 283, 195, and 47 genes, respectively. A total of 53 (C1), 209 (C2), 121 (C3), and 12 (C4) genes were positively selected exclusively in each population. Note, however, that the dN/dS ratios on populations A3 and C2 may be biased as they derive from a single observation (*Tara Ocean’s* station 65). For the species with potential populations defined by an FST=0.10 threshold, namely MAST-4B and MAST-4E, we observed 71 and 21 positively selected genes in populations B1 and B2, with 62 and 12 genes exclusive to them, respectively. Similarly, we observed 74 and 33 positively selected genes in MAST-4E’s populations E1 and E2, of which 63 and 22 were exclusive to each population, respectively.

Even though various selected genes in specific populations had matching orthologs in the eggNOG database, most annotations pertained to environmental sequences that lacked associated functions. The few genes that matched a reference with a known function were related to housekeeping functions or metabolisms, providing limited evidence regarding positively selected functions in specific populations (**Supplementary Dataset S3**).

All genes from the different MAST-4 species were clustered to detect broad patterns based on their dN/dS ratio variation across stations. For each MAST-4 species, genes were compelled into 50 clusters of variable size based on dN/dS similarity (**Figure 3**). Clusters were named following the CXSY formula, where X is the cluster number (1 – 50), and Y is the number of genes within the cluster. For each gene cluster and station, average dN/dS values were computed considering all the genes within the cluster. Gene clusters were subsequently grouped into bundles based on their dN/dS patterns across stations. Bundles displayed an array of average dN/dS values, from very low (ratio 0 – 0.2), low (0.2 – 0.5), and intermediate (0.5 – 0.8) across all samples to values close to 1 or greater either across all samples or in specific locations or potential populations (**Figure 3**). Overall, a total of 5 (MAST-4A), 5 (MAST-4B), 4 (MAST-4C), and 7 (MAST-4E) groups of clusters were defined (**Figure 3 Y-axis**).

**Figure 3.**
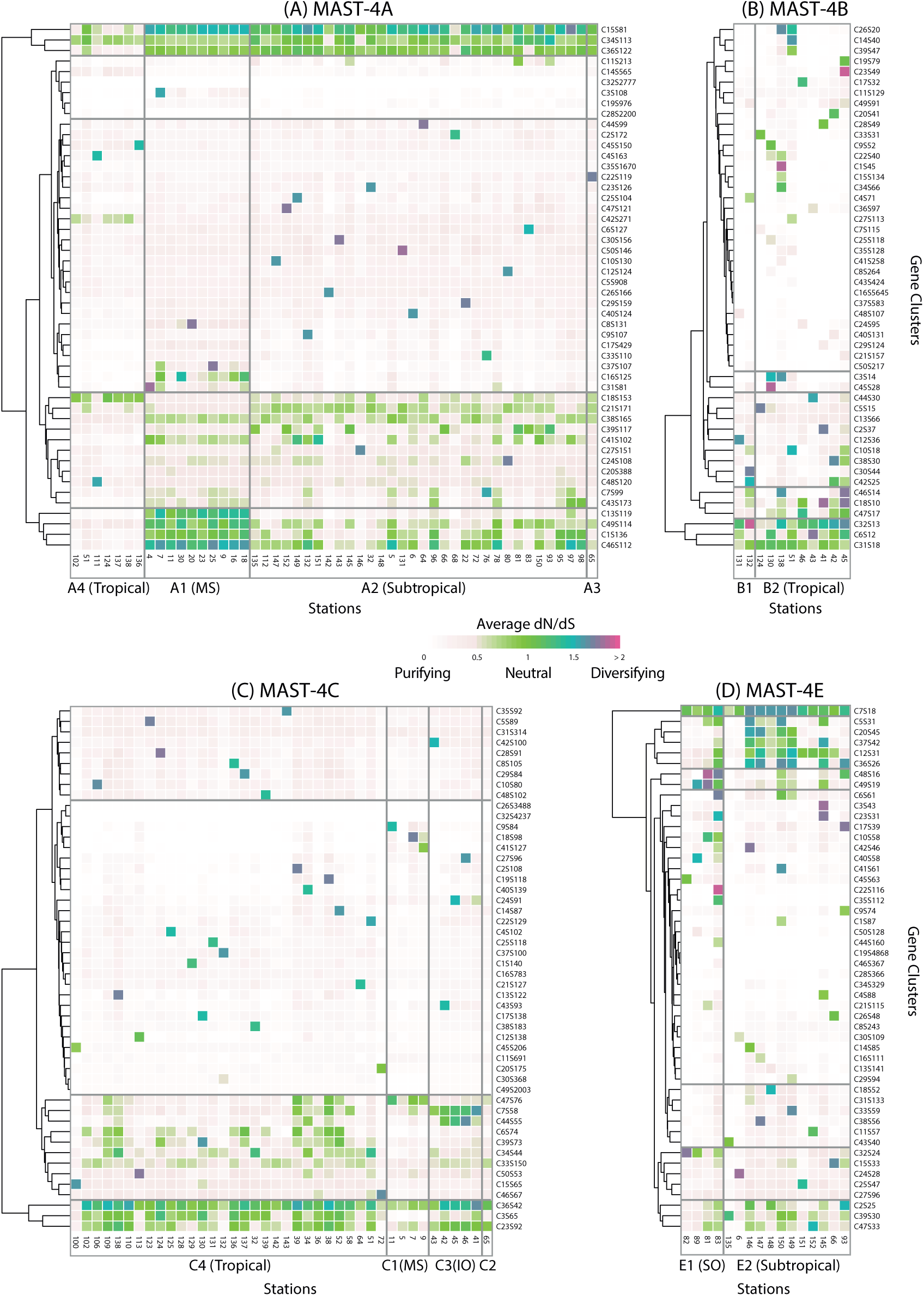
Average dN/dS of MAST-4 gene clusters across potential populations and *Tara Oceans* stations. Coding genes for each MAST-4 species (A [panel A], B [panel B], C [panel C], and E [panel D]) were clustered based on their similarities in dN/dS ratios across stations (UPGMA with “Manhattan” distance). The resulting dendrogram was cut at 50 clusters for each species to facilitate its representation. Colored tiles represent the clusters’ average dN/dS values per station. Darker colors represent a diversifying selection, while lighter colors represent a purifying selection. Gene cluster names are indicated as CXSY, where X is the cluster number (1 to 50), and Y is the number of genes within the cluster. Stations (X axes in each panel) are grouped based on potential populations (vertical lines). Note that the potential populations correspond to those defined in **Figure 2** based on FST thresholds (i.e., A1-4, B1-2, C1-4, E1-2). Specific putative populations have a tag indicating the ocean region they belong to (MS – Mediterranean, IO – Indian Ocean). Horizontal grey lines mark changes in average dN/dS ratios, separating the gene clusters into groups.

For each MAST-4 and gene cluster, MAST-4A featured, on average, the highest quantity of positively selected genes. Clusters C15S81, C34S113, and C36S122 had an average dN/dS > 1 in the putative populations A1 (Mediterranean Sea), A2 (Subtropical), and A3 (Subtropical - South-Africa coast), accounting for 316 genes (**Figure 3A**). Population A4 (Tropical) did not show clear patterns of positive selection in any of the gene clusters, except for specific stations, and several clusters displayed low dN/dS values, which may reflect the low abundance of this population. The Mediterranean Sea population (A1) appeared to have a clear positive selection for the gene clusters C13S119, C46S112, C49S114, and C1S136 (total of 481 genes) when compared to the other subtropical populations (A2 and A3) (**Figure 3A**). This suggests that these 481 genes, particularly those from C13S119, may represent the basis of the population A1 adaptive differentiation when compared against the reference MAST4-A genome. Large clusters of genes (e.g., C32S2777, C28S2200) showed evidence of purifying selection across all stations. They encompassed genes functionally annotated to general categories (**Figure S1 MAST-4A**), suggesting that these gene functions may be conserved throughout different oceanic regions.

Although MAST-4C had a similar number of predicted genes to MAST-4A, it exhibited, on average, a stronger purifying selection (dN/dS < 1; **Figure 3C**). This contrasts with the overall higher level of positive selection observed in species A, which had a greater number of gene clusters with a mean dN/dS > 1 (**Figure 3A**). In MAST-4C, clusters C26S3488, C32S4237, C11S691, and C49S2003 had a mean dN/dS close to 0 across all stations, accounting for 9,728 genes (64.1% of the total number of genes) (**Figure 3C**). Still, a few genetic clusters indicated positive selection. For example, clusters C36S42 and C23S92 (134 genes) showed positive selection across all delineated populations, particularly C2 (Subtropical), C3 (Indian Ocean), and C4 (Tropical) (**Figure 3C**). In comparison, positively selected gene clusters C7S58 and C44S55 (113 genes) were specific to the Indian Ocean (C3), while C47S76 was specific to the Mediterranean (C1) and the Red Sea (stations 38 and 39 from C4) (**Figure 3C**).

MAST-4B showed a more erratic distribution of positive selection across stations. Except for clusters C32S13, C6S12, and C31S18 (a total of 43 genes) that showed positive selection not associated with specific basins, the remaining 47 clusters displayed dispersed peaks of positive selection across stations (**Figure 3B**). Several other clusters displayed purifying selection across all or most stations (e.g., C16S5645), representing a substantial proportion of all the genes (**Figure 3B**). Lastly, MAST-4E showed dN/dS differences between specific North Atlantic stations (from station 145 to 150) and the rest of the stations. Genetic clusters C5S31, C20S45, C37S42, C12S31, and C36S26 displayed greater dN/dS values in the North Atlantic (**Figure 3D**). In contrast, only cluster C7S18 showed an overall positive selection across all analyzed stations (**Figure 3D**). As in the other MAST-4s, large clusters of genes (e.g., C19S4868) showed evidence of purifying selection across all stations, possibly pointing to conserved genes across populations (**Figure 3D**).

None of the clusters examined for species A, B, and E displayed an enrichment of specific eggNOG functions. Genes belonging to distinct functional categories were identified in all clusters (**Figure S1**). Additionally, except for a handful of gene clusters, they all had at least one gene with an unknown function. In contrast, for MAST-4C, the cluster C36S42 (positively selected in all potential populations) showed a larger proportion of genes within the *Chromatin structure and dynamics* category, while clusters C7S58 and C44S55 (positively selected in population C3) both displayed a larger proportion of genes within the *Carbohydrate transport and metabolism* category (**Figure S1 MAST-4C**). Nevertheless, certain functional patterns were discernible even without specific enrichment among species A, B, and E. The four MAST-4A gene clusters (*i.e.*, C13S119, C46S112, C49S114, and C1S136) that exhibited positive selection in the Mediterranean Sea (population A1) featured genes associated with *Replication, recombination and repair*, *Transcription and Translation, ribosomal structure, and biogenesis* categories (**Figure S1 MAST-4A**). MAST-4B’s most positively selected gene cluster, C32S13, comprised genes from the categories *Cell cycle control, cell division, chromosome partitioning,* and *Cytoskeleton* (**Figure S1 MAST-4B**). In turn, the cluster with the highest positive selection in MAST-4E, C7S18, displayed genes belonging to *Replication, recombination, and repair,* and *Amino acid and Inorganic ion transport and metabolisms* (**Figure S1 MAST-4E**).

## Discussion

The amount of genetic diversity in populations of uncultured marine microbes and how that diversity is structured over space and time remain largely unexplored. Here, we examined the population genomic diversity and differentiation of MAST-4 species, crucial unicellular predators across the surface global ocean. We observed differences in the number of variants per kilobase (Kb) in MAST-4 species, with an average of 87% being SNVs. The most abundant MAST-4 species, A and C, with larger and more complete genomes, exhibited higher variant numbers (18.2 and 14.0 variants per Kb of the genome, respectively) compared to the less abundant species, B and E, with smaller genomes (4.5 variants per Kb for both).

Although larger genomes provide more sites for potential mutations and a higher abundance of transposable elements, which can lead to more genetic variants^58,59^, microbial species with similar genome sizes have been reported to display different numbers of variants. For instance, the marine picoeukaryote *Bathycoccus prasinos*, with a genome size approximately half that of MAST-4B and E, showcased around 200,000 SNPs (13.3 SNVs per Kb) when analyzed in the same *Tara Oceans* dataset used in this work^17^. *Bathycoccus* is a relatively abundant taxon widely distributed in the ocean, similar to MAST-4 species A and C^26,27^. While few examples of genetic diversity are available for marine protists, a more substantial body of research exists for prokaryotes. For instance, the archaeon *Sulfolobus islandicus*, which has a genome size of about 2.6 Mb, exhibited around 8,100 SNVs (∼ 3.1 SNVs per kb)^60^ with a maximum FST similar to that of MAST-4A and C (maximum FST > 0.5), albeit it thrives in more restricted habitats like volcano springs^61^. However, this study did not use a metagenome-based population genomics approach but compared the genomes of 10 strains from one volcano spring sample^61^. Additionally, the highly abundant bacterium SAR11, with a smaller genome size (∼1.2 Mbp) than MAST-4^15^, revealed over one million genomic variants, specifically Single Amino Acid Variants (SAAVs), in 799 core genes for 21 population genomes over 103 metagenomic samples, of which 93 are from the *Tara Oceans* expedition (comparable geographic range as our study).

Abundance and distribution may be crucial factors in determining the number of genomic variants^11^. Greater abundance can increase genetic diversity by providing more opportunities for mutations and recombination events. As a result, abundant organisms are more likely to accumulate mutations because they undergo more replication events, providing more opportunities for mutations to occur purely by chance^62^. In turn, widespread organisms may have adapted to diverse environmental conditions, resulting in more genomic variants than organisms with a more limited habitat distribution^11^. Therefore, both abundance and distribution (or dispersal) may contribute to the genetic complexity of microbial populations and influence how these populations adapt to different niches. This is consistent with our findings, which show that the more abundant and widely distributed MAST-4A and C exhibited more nucleotide variants and higher population differentiation than the less abundant and more geographically restricted MAST-4B and E.

Nucleotide diversity, as a measure of the degree of polymorphism within a population, quantifies the average difference between all pairs of individuals at the nucleotide level. A low nucleotide diversity could be associated with population bottlenecks, selective sweeps, or rapid geographic or habitat expansion^63,64^. Conversely, a high nucleotide diversity may reflect a long history of diversification within a specific region or a prolonged period of high population abundance^63^. The higher nucleotide diversity observed in MAST-4A’s Mediterranean population A1 and MAST-4C’s C3 population in the Indian Ocean compared to the other populations could reflect a longer diversification and recombination history in those regions and higher historical abundances. Environmental conditions may have driven the emergence of regionally adapted populations that subsequently diversified, as evidenced by multiple positively selected genes in these populations (**Figure 3**). Specifically, population A1 experiences higher average salinity than other MAST-4A populations (due to its location in the Mediterranean Sea; **Table S2**). In turn, population C3 experiences water temperatures that are, on average, four °C higher than that experienced by other MAST-4C populations (**Table S2**). The comparatively low nucleotide diversity observed in other MAST-4A, B, C, and E populations could be linked to relatively recent geographic expansions of these populations. This is consistent with previous findings indicating early cladogenesis in MAST-4, with most current diversity represented by closely related taxa (low-rank diversity)^65^.

The population differentiation observed in MAST-4 species may be counterintuitive given the presumed high turnover rate of the genomic composition of the communities in the surface open ocean^66^. Previous studies using the 18S rDNA-ITS1 marker indicated spatial structuring in MAST-4A and E, suggesting sub-classes or populations influenced by seawater temperature^34^. Our genomic approach delineated populations by analyzing SNVs across the entire genome instead of relying on specific markers, which delineated more MAST-4 populations than those identified using the ITS1 marker in Rodríguez-Martínez *et al*.^34^ For instance, MAST-4A’s ITS1 region from different locations (Mediterranean Sea, Indian Ocean, North Atlantic Ocean, North Pacific Ocean, and the southern hemisphere) showcased little divergence with low FST values between them, forming one clade. In turn, our genomic approach suggests the presence of well-defined MAST-4A putative populations: one in the Mediterranean Sea, another in the open subtropical ocean, and a last in the tropical ocean. Moreover, the number of subclades delineated for MAST-4B, C, and E species based on the ITS1 marker is the same (two or more) as the number of putative populations defined by our genomic approach. However, Rodríguez-Martínez *et al*.^34^ found that the MAST-4B subclade was the most genetically differentiated of the group (highest FST), contrasting with our results in which, along with MAST-4E, it is the species with the lowest genomic differentiation. These differences suggest that the ITS and genome-wide SNVs capture distinct signals or dimensions of population differentiation and diversification.

Using UPGMA dendrograms based on FST distances and thresholds ranging between 0.15 and 0.10, we identified four potential populations for MAST-4A and C and two for MAST-4B and E. The FST index has been used to delineate genomic populations in marine organisms, from fish^13,67^ to microbes^15,68^, and values above 0.25 indicate high genetic differentiation^69^. Our analyses revealed two patterns based on the maximum FST divergence observed among the four MAST-4 species. Firstly, MAST-4A and C exhibited a substantial maximum divergence (maximum FST of 0.56 and 0.54, respectively), surpassing the threshold of 0.25 but falling slightly below that observed among allopatric populations of the diatom *Picea pungens* (maximum FST = 0.76), which experiences substantial limitations to gene flow^70^. Secondly, MAST-4B and E displayed moderate divergence (maximum FST of 0.14 and 0.21, respectively), akin to other cosmopolitan marine organisms without geographic constraints to gene flow, such as the diatom *Thalassiosira rotula* (maximum FST = 0.14)^71^. Despite the limited information, the FST values we obtained for the four MAST-4 species appear consistent with those reported for other marine protists.

Our analyses indicated that MAST-4A and C (featuring the highest FST) have undergone more extensive population differentiation than MAST-4B and E (with lower FST). The emergence of population differentiation in the open ocean and semi-enclosed seas, like the Mediterranean Sea, may be attributed to restricted gene flow or adaptation to specific environmental conditions. Known factors, such as geographic distance, can act as barriers, limiting the dispersal of marine plankton and consequently promoting population divergence in MAST-4^68^. Despite the assumed high-dispersal capability of MAST-4 in the open ocean^23,72^, recent studies underscore the substantial impact of dispersal limitation^73^ on global ocean surface picoplankton. Moreover, local adaptation to different niches could contribute to population differentiation in MAST-4^27^, exemplified by populations A1 and C3, which experience different salinity and temperature. This is supported by the multiple positively selected genes detected in these populations (**Figure 3**).

Temperature and salinity have been reported as influential variables in structuring microbial populations in the ocean. For example, salinity is a primary driver of the population structure of the Baltic Sea bacterioplankton^52^, and temperature was significantly correlated with *Prochlorococcus* ecotype abundances in the Atlantic Ocean^74^. Similarly, SAR11 strains are adapted to different current temperatures^15^. Comparable patterns have been observed in microbial eukaryotes. For example, *Bathycoccus prasinos* appears to be adapted to temperature variations across different depths in the North Atlantic^17^. Our study showcases a significant correlation between population-level genomic divergence in MAST-4 and temperature and salinity across the surface global ocean, emphasizing these variables as possible critical factors shaping MAST-4 population structure.

Previous studies have identified temperature as the primary factor shaping the biogeography of MAST-4 and other MAST lineages in the open ocean^27,28,72^. However, the extent of temperature’s influence on the population structure of MAST-4 remained unclear until now. Collectively, both marker-specific^34^ and genome-wide (this study) approaches underscore the critical role of temperature in shaping population structure within the MAST-4. Both methodologies predict that at least one MAST-4E subclade and population (E1, Southern Ocean) are adapted to cold waters. Here, we also observe a well-defined putative population of MAST-4C (C3, Indian Ocean) populating warmer waters and featuring several positively selected genes. Additionally, Rodríguez-Martínez *et al*.^34^ showcased that salinity might be one of the fundamental abiotic factors influencing population structure within temperature-defined groups. This agrees with the genomic divergence observed in our MAST-4A populations, where the MAST-4A Mediterranean A1 and Subtropical A2 populations have a difference of less than one °C for their average temperatures (20.9 and 20.2 °C respectively) and a significant difference in salinity of ca. 2 PSU (**Table S2**). Populations A1 and A2 are differentiated with distinct positively selected genes (**Figure 3**). However, it remains unclear whether this phenomenon reflects direct adaptation to salinity, given that the 2 PSU difference observed in marine samples represents a relatively narrow range of variation. An alternative scenario is that this differentiation reflects adaptation to different water masses, such as those in the Mediterranean Sea^75^.

In Latorre *et al.*^27^, we examined how adaptation to diverse ecological conditions has driven evolutionary diversification among MAST-4 species. Adaptive evolution is apparent in the genetic variation observed across the MAST-4 populations in this study, highlighting the roles of temperature and salinity in shaping their evolutionary trajectories at finer scales. Building on the recently proposed evolutionary history of MAST-4^27^, the divergence of MAST-4A and MAST-4C into distinct ecological niches is further amplified by the emergence of separate populations within these species. Specifically, MAST-4A, initially adapting to temperate waters to avoid competition with MAST-4C, may have experienced further differentiation within its geographic range. Two main populations of MAST-4A seem to have emerged: one adapted to the Mediterranean Sea and another inhabiting the global ocean’s subtropical waters. In turn, MAST-4C, which appears to be ancestral to warm waters^27^, has further diversified into two main warm-water populations: C3, restricted to the warmer waters of the Indian Ocean, and C4, found in other tropical regions. Several positively selected genes identified in C3 may be associated with adaptations to higher temperatures (**Figure 3**). Overall, the evolutionary diversification of MAST-4 exemplifies a key ecological principle: lineages may emerge by adapting to different niches, and subsequent adaptive evolution within these niches can lead to distinct populations specialized for even more specific microenvironments. Although this is predicted by evolutionary theory, few examples are available for free-living microbes.

To better understand the adaptive mechanisms in MAST-4 populations, we examined the ratios of non-synonymous to synonymous mutations for all protein-coding genes in each species. Our focus was mainly on those genes that showed evidence of diversifying selection (dN/dS > 1) across potential populations or ocean regions. This analysis identified positively selected genes that formed gene clusters with similar dN/dS patterns, corresponding to specific populations or oceanic basins within the MAST-4 species in several cases. Some gene clusters showed positive selection across all samples, suggesting that these genes may play a key role in the genomic differentiation of several populations. We also identified gene clusters with distinct dN/dS patterns specific to certain populations. For example, in MAST-4A, gene clusters in the Mediterranean population (A1), characterized by higher salinity, exhibited higher average dN/dS ratios than those in other stations (**Figure 3A**). Similarly, the C3 population of MAST-4C showed gene clusters with higher dN/dS ratios at stations in the Indian Ocean with elevated temperatures compared to other stations (**Figure 3C**). This may indicate adaptations to higher salinity or temperature found in the Mediterranean Sea and Indian Ocean, respectively, compared to other tropical and subtropical waters (**Table S2**).

These findings are consistent with other studies that have applied genomic approaches to marine microorganisms. For instance, Delmont and Eren^76^ applied metapangenomic analyses to *Prochlorococcus* across the global ocean, identifying a small number of gene clusters that might be linked to differences in fitness among closely related members. These clusters, defined by the presence or absence of accessory genes, play a crucial role in adaptation to different oceanic regions. In contrast, the gene clusters we identified in MAST-4 are delineated by patterns of positive selection rather than accessory gene variation. This distinction might indicate that MAST-4 populations may rely on different mechanisms of regional adaptation driven by evolutionary changes in specific genes. Delmont et al. (2019)^15^ also employed a metagenomic approach based on single-amino acid variants (SAVs) to examine evolutionary processes within *SAR11*. This study revealed that specific SAVs were associated with environmental variables like temperature and nutrient availability, demonstrating how fine-scale genetic variants can drive microbial adaptation across diverse environments.

The difference in dN/dS patterns across specific gene clusters and potential populations could indicate the role of these genes in conferring adaptive advantages in response to local environmental conditions. This aligns with findings in various studies where key pathways were involved in temperature adaptation. For example, genes involved in the *replication, transcription, and translation* of DNA and RNA; *carbohydrate transport and metabolism;* and *amino acid and inorganic ion transport and metabolism* are among the functional categories regulated by temperature in the bacteria *Sphingopyxis alaskensis*^77^. These categories were also found in gene clusters with high dN/dS in MAST-4 species and could play a similar role. By applying a similar genome-centric approach to bacterial Metagenome-Assembled Genomes (MAGs) from the Baltic Sea, it was found that genes encoding for core functions likely experienced higher purifying selection than others^52^. Other genes associated with secondary metabolisms, such as anti-microbial resistance genes, displayed the highest dN/dS values, pointing to targeted diversifying selection^52^. In all four MAST-4 species, several large gene clusters containing both core and secondary genes exhibited purifying or neutral selection across most stations. Clusters with high dN/dS ratios, indicating positive selection, also included core and secondary genes. This suggests that population adaptation in MAST-4 may be distributed across both core and secondary genes.

In protists, mutations associated with cold adaptation have been identified in a protein transporter involved in exporting cations in *B. prasinos*^17^. MAST-4 species could similarly thrive in colder water environments through comparable adaptations. For instance, protein transporters involved within the *Replication, recombination, and repair,* and *Amino acid and Inorganic ion transport and metabolisms* categories were found positively selected in MAST-4E, which suggests population adaptations to cold water (*e.g.*, population E1, present in cold South Atlantic waters). Interestingly, one of the key proteins involved in cold adaptation in *B. prasinos*, the eEF3 protein^17^, was found to be exclusively under positive selection in the MAST-4A1 population (COG0488; **Supplementary Dataset S3**). This finding suggests that the eEF3 protein may play a role in adapting MAST-4A populations to diverse environmental conditions. However, whether this adaptation is primarily driven by temperature or other environmental factors remains unclear.

A few functional categories identified as positively selected in MAST-4 populations align with genes found to be up and down-regulated under different temperature conditions in transcriptomic studies of diatoms^78^. This overlap suggests a potential link between gene expression plasticity and evolutionary adaptation in microbial eukaryotes facing varying thermal environments. For example, positively selected genes related to *Replication*, *Transcription* and *Translation, ribosomal structure, and biogenesis* in MAST-4A1 are similarly enriched in cold-diatom species. However, despite this similarity, our results do not indicate temperature adaptation in MAST-4A1 compared to the other subtropical populations. Instead, the pattern of positive selection suggests that these genes might be crucial for adaptation to other environmental factors rather than directly to temperature.

Similarly, positively selected genes involved in *Amino acid transport and metabolism* in MAST-4E populations are among those enriched functions in cold-adapted diatom species, where they may mitigate the effects of lower translation efficiency at cooler temperatures. Additionally, positively selected genes in the *Carbohydrate transport and metabolism* category in MAST-4C3 and *Cell cycle-*related categories in MAST-4B populations were identified as crucial to temperature adaptation in diatoms. These findings suggest that pathways subject to temperature-responsive gene regulation in diatoms may also be under evolutionary selective pressures in MAST-4 populations, as evidenced by the observed dN/dS ratios. Gene expression plasticity likely enables microbial eukaryotes to respond rapidly to environmental stress, while natural selection refines these adaptations over time. This dynamic interplay offers valuable insights into how microbial eukaryotes adapt and thrive across the diverse and changing environments of the global ocean.

Although our focus is on metagenomic data, the insights from transcriptomics underscore the dynamic nature of gene expression in response to environmental changes. A dN/dS > 1 points to nucleotide changes that can only alter the protein’s final function, expression, or tertiary structure^79^ on coding sequences. Yet, approximately 80% of MAST-4’s variants were located in non-transcribed regions, which could also influence gene expression^80^. Furthermore, despite not altering the protein sequence in yeasts, synonymous mutations can impact mRNA expression and fitness^81,82^. This suggests that genes with a relatively low dN/dS ratio may still be significant for population adaptation. Therefore, given the relevance of differential gene expression in response to environmental changes, particularly temperature, in other microorganisms^83,84^, it is plausible that non-coding and synonymous mutations that modify gene expression may also contribute to shaping the population structure of MAST-4 in the surface global ocean.

Our findings provide new insights into the population genomics of MAST-4, a key unicellular predator in the global ocean. MAST-4A exhibits high genetic divergence, with at least four potential populations adapted to different temperatures and possibly other environmental conditions. In contrast, MAST-4B shows moderate genetic divergence, with its population structure strongly influenced by temperature, particularly in tropical regions. MAST-4C, which predominates in tropical waters, also displays genetic differentiation driven by temperature; however, unlike MAST-4B, it exhibits highly divergent potential populations. Lastly, MAST-4E demonstrates moderate genetic divergence but provides evidence of potential populations adapted to varying temperature ranges, spanning subtropical to subpolar regions. MAST-4 represents a group of uncultured heterotrophic flagellates, with species characterized by well-defined populations that appear to have evolved through niche adaptation. Understanding these populations at a finer scale contributes to our knowledge of marine food webs and their resilience to global change.

## Acknowledgments

We thank all *Tara Oceans* 2009-2013 expedition scientists and the Single Cell Genomics Center (Bigelow Laboratory) staff for generating single amplified genomes. Bioinformatics analyses were performed at the MARBITS platform of the Institut de Ciències del Mar (ICM) [https://marbits.icm.csic.es]. We thank Pablo Sánchez and Lidia Montiel for their assistance with bioinformatics. FL was supported by the Spanish National Program FPI 2016 (BES-2016-076317, MICINN, Spain). RL was supported by a Ramón y Cajal fellowship (RYC-2013-12554, MINECO, Spain). RM was supported by EPIC (PID2022-137508NB-I00, MICINN). This work was supported by the project INTERACTOMICS (CTM2015-69936-P, MINECO, Spain), MicroEcoSystems (240904, RCN, Norway), and MINIME (PID2019-105775RB-I00, MICINN, Spain) to RL.

## SUPPLEMENTARY MATERIAL

### Supplementary Figures

**Figure S1.**
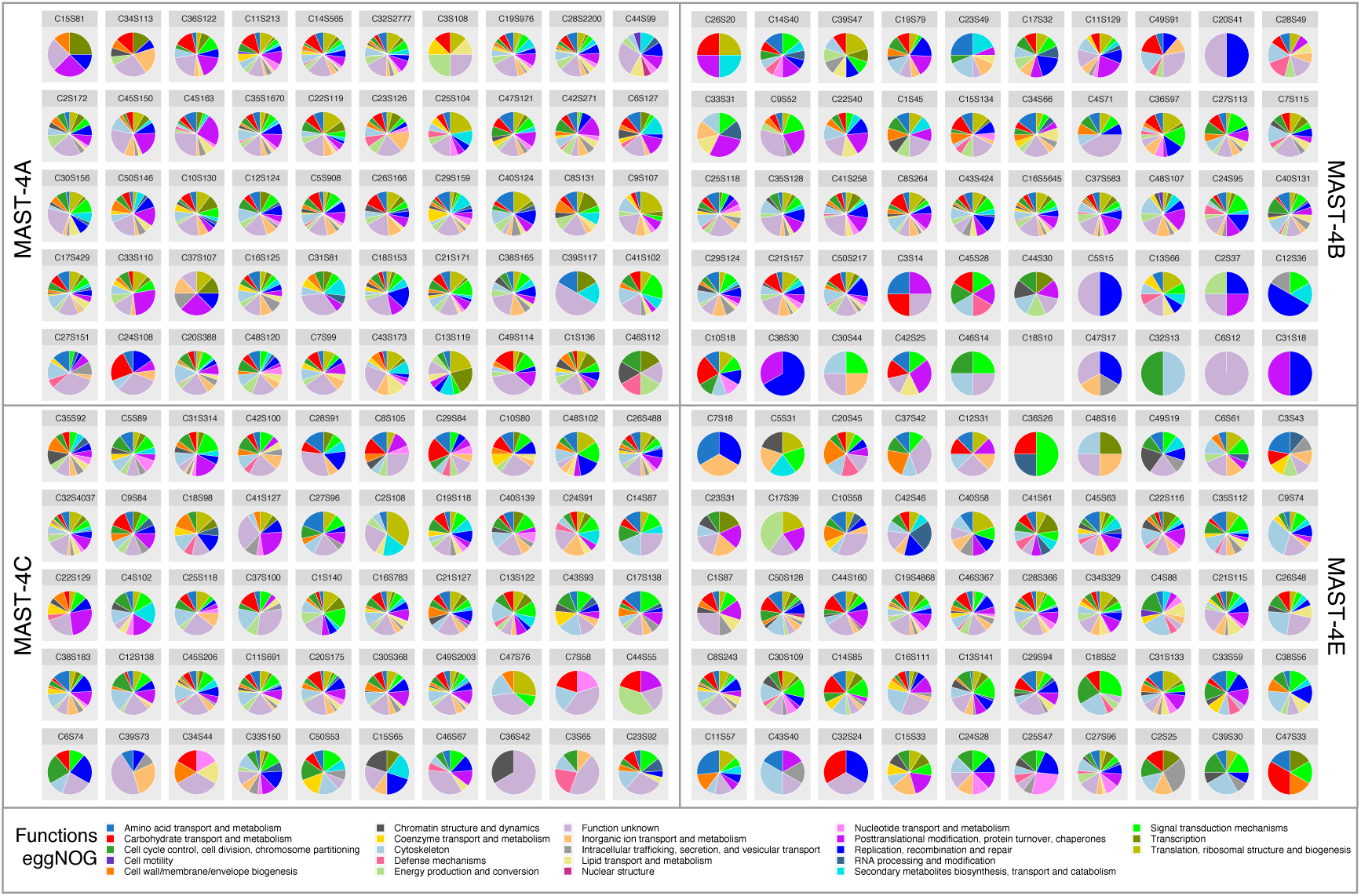
Functional annotation of genetic clusters for MAST-4 species using the eggNOG database. A total of 50 gene clusters were delineated based on similarities of dN/dS ratios across stations (see **Figure 3**). Gene cluster names are CXSY, where X is the cluster number (1 to 50), and Y is the number of genes within the cluster. Genes without a hit in the database were not considered. A cluster without a pie chart indicates that no gene was found in the eggNOG database.

### Supplementary Tables

**Table S1.** Number of variants and their effects by type for each MAST-4 species across all stations, both in counts and percentage.

|  | MAST-4A |  | MAST-4B |  | MAST-4C |  | MAST-4E |  |
| --- | --- | --- | --- | --- | --- | --- | --- | --- |
|  | N° | % | N° | % | N° | % | N° | % |
| *Number of variants by type* |  |  |  |  |  |  |  |  |
| **SNVs** | 735,967 | 85.18 | 115,191 | 87.87 | 585,004 | 87.50 | 120,952 | 88.06 |
| **MNVs** | 62,463 | 7.23 | 10,755 | 8.20 | 60,579 | 9.06 | 8,317 | 6.06 |
| **INS** | 29,676 | 3.43 | 2,066 | 1.58 | 9,898 | 1.48 | 3,148 | 2.29 |
| **DEL** | 26,765 | 3.10 | 2,545 | 1.94 | 10,843 | 1.62 | 4,277 | 3.11 |
| **MIXED** | 9,138 | 1.06 | 534 | 0.41 | 2,289 | 0.34 | 663 | 0.48 |
| *Number of effects by impact* |  |  |  |  |  |  |  |  |
| **HIGH** | 3,778 | 0.14 | 353 | 0.08 | 1,463 | 0.07 | 439 | 0.10 |
| **LOW** | 287,944 | 10.89 | 61,316 | 14.00 | 333,785 | 15.20 | 38,496 | 8.62 |
| **MODERATE** | 273,784 | 10.36 | 32,464 | 7.41 | 173,654 | 7.91 | 32,342 | 7.24 |
| **MODIFIER** | 2,078,495 | 78.61 | 343,797 | 78.51 | 1,687,836 | 76.83 | 375,597 | 84.05 |
| *Number of effects by functional class* |  |  |  |  |  |  |  |  |
| **MISSENSE** | 238,756 | 46.05 | 27,518 | 32.17 | 145,238 | 31.73 | 29,076 | 43.86 |
| **NONSENSE** | 765 | 0.15 | 79 | 0.09 | 316 | 0.07 | 50 | 0.08 |
| **SILENT** | 279,002 | 53.81 | 57,953 | 67.74 | 312,180 | 68.20 | 37,172 | 56.07 |
| *Number of effects by region* |  |  |  |  |  |  |  |  |
| **DOWNSTREAM** | 932,929 | 35.29 | 164,630 | 37.59 | 817,223 | 37.20 | 167,049 | 37.38 |
| **EXON** | 562,897 | 21.29 | 93,864 | 21.43 | 507,454 | 23.10 | 70,884 | 15.86 |
| **INTERGENIC** | 264,479 | 10.00 | 33,604 | 7.67 | 141,793 | 6.46 | 59,028 | 13.21 |
| **INTRON** | 34,230 | 1.30 | 3,360 | 0.77 | 17,960 | 0.82 | 7,061 | 1.58 |
| **SPLICE_ACCEPTOR** | 307 | 0.01 | 32 | 0.01 | 136 | 0.01 | 39 | 0.01 |
| **SPLICE_DONOR** | 307 | 0.01 | 28 | 0.01 | 155 | 0.01 | 45 | 0.01 |
| **SPLICE_REGION** | 1,995 | 0.08 | 209 | 0.05 | 1,157 | 0.05 | 309 | 0.07 |
| **TRANSCRIPT** | 2 | 0.00 | 1 | 0.00 | 3 | 0.00 | 0 | 0.00 |
| **UPSTREAM** | 846,839 | 32.03 | 142,199 | 32.47 | 710,840 | 32.36 | 142,457 | 31.88 |
| **UTR_5_PRIME** | 16 | 0.00 | 3 | 0.00 | 17 | 0.00 | 2 | 0.00 |

SNV – Single-Nucleotide Variant; MNVs – Multiple-Nucleotide Variants; INS – Insertions; DEL – Deletions; MIXED – Mix of Multiple-Nucleotide Variants and INDELS. Impact categories: HIGH – disruptive impact on the protein (truncation or loss of function); MODERATE – potential change in the protein effectiveness; LOW – unlikely to change protein functionality; and MODIFIER – for non-coding variants or variants for which it is difficult to predict impact. Functional class categories: MISSENSE – variants that change the amino acid; NONSENSE – variants that add a STOP codon (truncate the protein); and SILENT – variants that do not change the amino acid. Region categories: DOWNSTREAM – downstream of a gene; EXON – variant hits an exon; INTERGENIC – variant in an intergenic region; INTRON – variant hits an intron; SPLICE\_ACCEPTOR – variant hits a splice acceptor site (defined as two bases before exon start, except for the first exon); SPLICE\_DONOR – variant hits a splice donor site (defined as two bases after coding exon end, except for the last exon); SPLICE\_REGION – variant in which a change has occurred within the region of the splice site, either within 1-3 bases of the exon or 3-8 bases of the intron; TRANSCRIPT – variant hits a transcript; UPSTREAM – upstream of a gene; and UTR\_5\_PRIME – variant hits 5'UTR region.

**Table S2.** Potential populations for each MAST-4 species. Average temperature (°C) and salinity (PSU) values are listed for each potential population with its standard deviation. See **Figure 2**.

| Species | Potential population | Mean Temperature (°C) | Mean Salinity (PSU) | Region | n° Stations |
| --- | --- | --- | --- | --- | --- |
| MAST-4A | A1 | 20.9 ± 2.3 | 38.0 ± 2.3 | Mediterranean Sea | 10 |
|  | A2 | 20.2 ± 4.5 | 36.0 ± 4.5 | Subtropical | 32 |
|  | A3 | 21.8 | 35.4 | South Africa | 1 |
|  | A4 | 25.6 ± 1.6 | 34.7 ± 1.6 | Tropical | 7 |
| MAST-4B | B1 | 25.7 ± 0.8 | 35.2 ± 0.8 | Subtropical | 2 |
|  | B2 | 28.7 ± 1.6 | 34.8 ± 1.6 | Tropical | 9 |
| MAST-4C | C1 | 23.0 ± 2.0 | 37.5 ± 2.0 | Mediterranean Sea | 4 |
|  | C2 | 21.8 | 35.4 | South Africa | 1 |
|  | C3 | 30.0 ± 0.5 | 35.0 ± 0.5 | Indian Ocean | 5 |
|  | C4 | 26.0 ± 1.4 | 35.4 ± 1.4 | Tropical | 30 |
| MAST-4E | E1 | 8.5 ± 3.5 | 34.0 ± 3.5 | Sub-polar | 4 |
|  | E2 | 17.6 ± 2.1 | 35.8 ± 2.1 | Subtropical | 12 |

### Supplementary Datasets

**Supplementary Dataset S1.** Metagenomic samples from the *Tara Oceans* expedition.

**Supplementary Dataset S2.** Environmental metadata for *Tara Oceans* stations.

**Supplementary Dataset S3.** Functional annotation of genes exclusive to each MAST-4B genomic population, based on eggNOG and CAZy databases. Exclusively identified genes without functional annotation are not included. NOTE: Mean dN/dS values > 2 are annotated in the table as 2.

**Supplementary Dataset S4.** Functional annotation of genes exclusive to each MAST-4B genomic population, based on eggNOG and CAZy databases. Exclusively identified genes without functional annotation are not included.

**Supplementary Dataset S5.** Functional annotation of genes exclusive to each MAST-4C genomic population, based on eggNOG and CAZy databases. Exclusively identified genes without functional annotation are not included.

**Supplementary Dataset S6.** Functional annotation of genes exclusive to each MAST-4E genomic population, based on eggNOG and CAZy databases. Exclusively identified genes without functional annotation are not included.

## Notes

https://github.com/franlat/mast4_dnds

### Competing Interest Statement

The authors have declared no competing interest.

